# Understanding Ecological Systems Using Knowledge Graphs: An Application to Highly Pathogenic Avian Influenza

**DOI:** 10.1101/2024.09.05.611483

**Authors:** Hailey Robertson, Barbara A. Han, Adrian A. Castellanos, David Rosado, Guppy Stott, Ryan Zimmerman, John M. Drake, Ellie Graeden

**Affiliations:** Yale School of Public Health, Department of Epidemiology of Microbial Diseases, New Haven, CT 06510; Center for Global Health Science and Security, Georgetown University, Washington DC 20007; Massive Data Institute and Center for Global Health Science and Security, Georgetown University, Washington DC 20007; Cary Institute of Ecosystem Studies, Box AB Millbrook, NY 12545; Institute of Bioinformatics, Department of Infectious Diseases, Department of Epidemiology and Biostatistics, Center for Ecology of Infectious Diseases, Center for Applied Pathogen Epidemiology and Outbreak Response, University of Georgia, Athens, GA 30602; Odum School of Ecology and Center for the Ecology of Infectious Diseases, University of Georgia

## Abstract

Ecological systems are complex. Representing heterogeneous knowledge about ecological systems is a pervasive challenge because data are generated from many subdisciplines, exist in disparate sources, and only capture a subset of important interactions underpinning system structure, resilience, and dynamics. Knowledge graphs have been successfully applied to organize heterogeneous data systematically and to predict new linkages representing unobserved relationships in complex systems. Though not previously applied broadly in ecology, knowledge graphs have much to offer in an era of global change when system dynamics are responding to rapid changes across multiple scales simultaneously. We developed a knowledge graph to demonstrate the method’s utility for ecological problems focused on highly pathogenic avian influenza (HPAI), a highly transmissible virus with a broad animal host range, wide geographic distribution, and rapid evolution with pandemic potential. We describe the development of a graph to include a wide range of data related to HPAI including pathogen-host associations, animal species distributions, and human population demographics, using a semantic ontology that defines relationships within the data and between datasets. We use the graph to perform a set of proof-of-concept analyses validating the method and identifying new relationships and features of HPAI ecology, underscoring the generalizable value of knowledge graphs to ecology including their utility in revealing previously known relationships between entities and generating testable hypotheses in support of a deeper mechanistic understanding of ecological systems.

## Introduction

The complexity of ecological systems is well-appreciated (Solé & Levin, 2022). However, analysis in the field has largely relied on pairwise or targeted integration of specific subfields that hinges on previously established foundational knowledge about relationships between the topical domains (Silk et al., 2022). For example, the population dynamics of key species like pollinators and predators and the outbreak and pandemic spread of novel pathogens are examples of phenomena that emerge from the confluence of multiple simultaneous processes and non-linear relationships (Brett et al., 2017; Sánchez-Garduño et al., 2014; Tan et al., 2022). Within disease ecology specifically, there has been a move to include social science research, including policy analysis and survey data about human behavior such as vaccine hesitancy, with the more traditional quantitative and biological analyses performed by ecologists. To facilitate this interdisciplinary work into disease ecology, however, we need to operationalize new methods to integrate data from these systems and models across domains (Friant, 2024; B. A. Han & Drake, 2016; Rivers et al., 2019; Todman et al., 2023; Woldehanna & Zimicki, 2015)

To successfully understand the relationships between these disparate systems, we need scalable methods to integrate broadly heterogeneous data and modeling methods. For example, remote sensing data from earth systems science can be combined with animal observation data to generate a composite understanding of phenology, abundance, migration, or extinction. Such synthetic analyses are increasingly common and offer a powerful deductive tool with which to better understand many dynamic processes in zoonotic disease outbreaks that are difficult to assess due in large part to data incompleteness.

Knowledge graphs (KGs) are graph-based structures that support data integration by encoding data points (nodes) and the relationships between them (edges), using semantic mappings to connect disparate data with shared underlying elements across domains and scales while avoiding taxonomic and unit misalignment. KGs are particularly effective in linking and structuring heterogeneous data to make them interoperable (i.e., connecting knowledge represented differently by different disciplines).

While not yet widely applied in ecology, the value of KGs has been demonstrated in various fields to explore interconnections between entities and detect anomalous patterns. For instance, biomedical KGs have been used for drug target discovery (Mohamed et al., 2019), prediction of potential drug-drug interactions (Ye et al., 2021), precision medicine (Chandak et al., 2023), and meta-analysis of diseases like COVID-19 (Pestryakova et al., 2022). Outside of biomedicine, KGs have integrated multi-source spatiotemporal data for natural disaster early warning systems and risk classification (Ge et al., 2022; Z. Yang et al., 2022), humanitarian response efforts (W. Li et al., 2023), life sciences (Callahan et al., 2024) and environmental research (F. Han et al., 2022; Zárate et al., 2019). The expanding applications of KGs, coupled with recent advancements in machine-learning models, open up new potential for sophisticated analysis of heterogeneous datasets across disciplines and scales.

Here, we describe the application of KG methods to an ecological system and demonstrate the utility of the graph in ecology with a case study on “bird flu” (highly pathogenic avian influenza A virus or HPAI) to demonstrate how an ontology semantically connects data from heterogeneous sources to create a knowledge graph that can support statistical modeling and analysis of heterogeneous data about this system.

### Building a knowledge graph for ecological analysis of highly pathogenic avian influenza

HPAI virus, specifically H5 and H7 subtypes, is an evolving threat to poultry, wildlife populations, and human health. In early 2020, an H5N1 lineage experienced a major resurgence and replaced the dominant H5N8 virus, spreading rapidly to all continents except for Australia (Xie et al., 2023). Since then, the H5N1 outbreak has caused unprecedented mass mortality in species that have never been detected with bird flu before – including many mammals (Graziosi et al., 2024). There is also evidence of significant mammal-to-mammal transmission, including among dairy cattle and detectable within milk, which has raised concerns about the rapid reassortment of the virus and its potential to cause a human pandemic (European Food Safety Authority (EFSA) et al., 2024; Petersen et al., 2024). Despite the virus’ zoonotic potential, relatively little is known about the dynamics of HPAI globally, such as how its viral evolution may impact geographic spread, host range, and the species-level attributes associated with infection (Runstadler & Puryear, 2024). Therefore, we use the case of HPAI (Influenza A/H5N1) to demonstrate how KGs can support rapid outbreak analysis in a fragmented data environment, highlight knowledge gaps, and support hypothesis generation about cross-species transmission and future spread.

## Methods

Constructing and applying a knowledge graph to support scientific inquiry involves five steps (Figure 1):

1. Identify data sources to populate the graph
2. Develop an ontology that represents the real-world relationships that you are trying to describe and connects them with each of the chosen data sources
3. Extract, transform, and load data using graph database software to construct the knowledge graph
4. Embed the knowledge graph to test its coherence and train models on the structure and patterns of the graph
5. Perform downstream modeling and analysis

**Figure 1.**
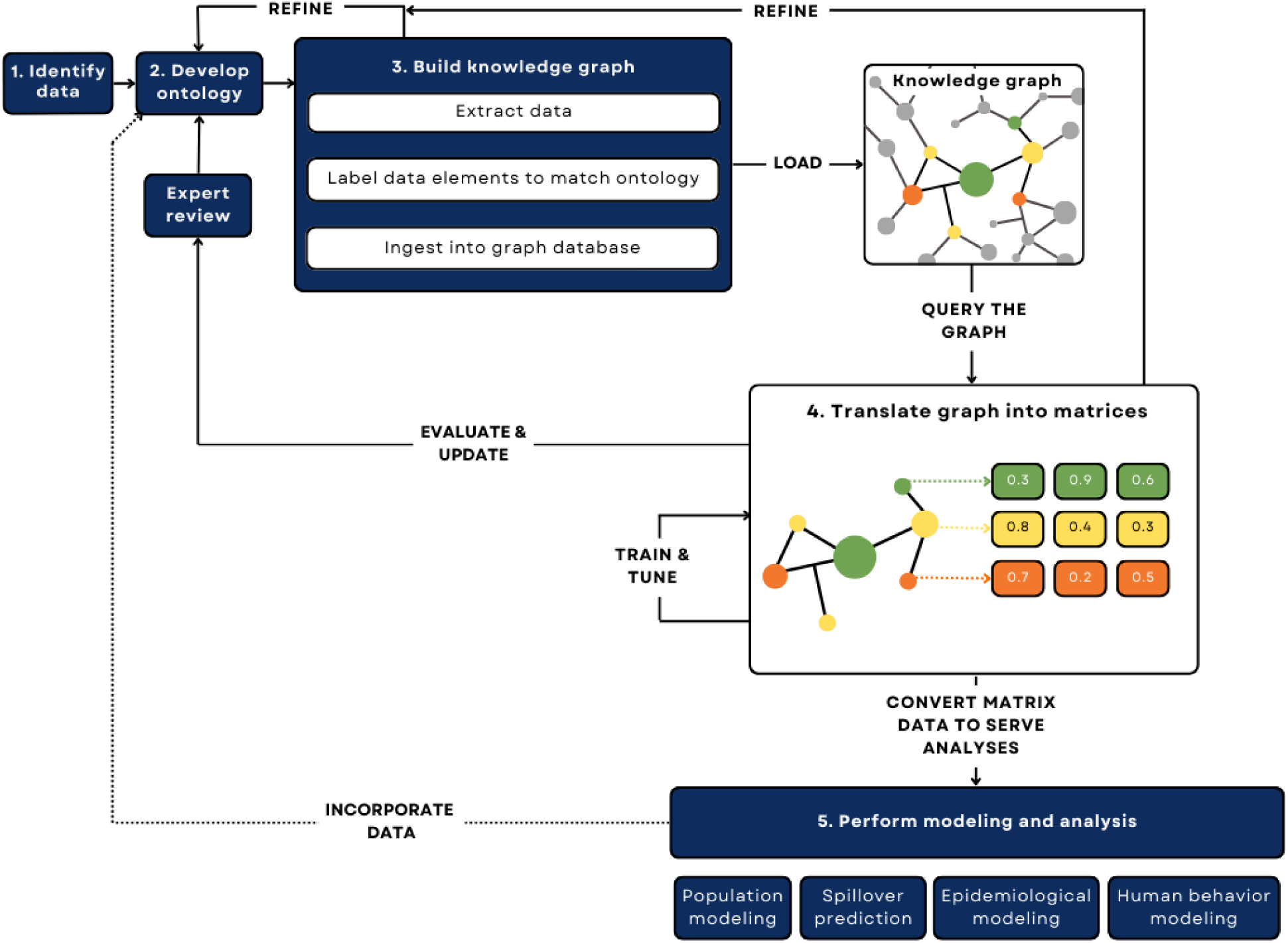
Process map depicting the steps to construct and use a custom knowledge graph from data identification to downstream machine learning tasks. 1) Identify data requirements based on research questions, inventory data sources, and define shared data elements. 2) Develop an ontology by defining nodes as a combination of one or more shared data elements, defining edges as relationships between nodes, and using attributes to describe nodes or edges with additional detail. 3) Build the graph by extracting data from each source, labeling data elements from each source to match the ontology as a node, edge, or attribute, and ingesting data into the graph database. Note: this describes the extract-transform-load (ETL) pipeline. 4) Translate the graph into matrices for analyses, such as routine statistics or machine learning models using knowledge graph embedding models or graph neural networks. 5) Perform analysis and modeling using the graph by converting the matrix data into parameters for population modeling, spillover prediction, epidemiological modeling, or human behavior modeling. Incorporate new, inferred knowledge into the graph if desired, refining the ontology and repeating the process as necessary.

### 1. Identify data

To understand zoonotic disease system dynamics and model human spillover risk, we first defined key information requirements and compiled corresponding datasets with information about species traits, population demographics, pathogen characteristics, environmental factors, and human behavior into a data inventory. Of the data in the inventoried datasets, we identified six classes: reports, events, populations, taxa, geographies, and samples (Suchanek et al., 2019). All datasets contained at least one, and many had three or more of these classes represented, typically represented as columns or properties that were used as the basis for semantic classification.

Although all 28 datasets were used to inform the shared data elements and ontology (described in the following section), we prioritized 7 key data sources to construct the initial graph given their relevance to zoonotic disease risk and shared concepts (e.g., classes) despite different taxonomies and structures. These were GeoNames (*GeoNames*, n.d.), NCBI Taxonomy (*Home - Taxonomy - NCBI*, n.d.), the Global Mammal-Parasite Database 2.0. (GMPD2) (Stephens et al., 2017), World Health Organization (WHO) FluNet (Flahault et al., 1998), World Organization for Animal Health (WOAH, formerly OIE) World Animal Health Information System (WAHIS) (*World Animal Health Information System*, 2021), United Nations (UN) World Population Prospects (WPP) (*World Population Prospects - Population Division - United Nations*, 2023), and the Coalesced Mammal Database of Intrinsic and Extrinsic Traits (COMBINE, (Soria et al., 2021).

GeoNames and NCBI Taxonomy were chosen for their stable unique identifiers (GeonamesId and TaxIds, respectively). These sources each have their own established ontologies and have been widely used to support knowledge graph and database development (Callahan et al., 2024; J. Lin et al., 2021; Wardeh et al., 2015).

GMPD2 provides data on the range of host-pathogen interactions for HPAI A(H5N1) and other pathogens (Stephens et al., 2017). While datasets like CLOVER, EID2, HP3, and Shaw (2020) offer wider species coverage (Gibb et al., 2021), GMPD2 uniquely provides public data downloads along with diagnostic details and coordinates for each host-pathogen association. We prioritized a host-pathogen dataset with geographic coverage to counter the assumption that a host-pathogen association observed in one population applied universally to the species.

FluNet (Flahault et al., 1998) was used to obtain comprehensive weekly reports on human flu occurrences at the country level. WAHIS provides animal case data for 85 globally notifiable pathogens, including extensive reporting on HPAI (S.-Y. Lin et al., 2023; *Terrestrial Code Online Access*, 2021). The data are updated in real time and accessible via a web portal (*WAHIS*, n.d.). Use of data from the WAHIS platform requires the following statement: “WOAH bears no responsibility for the integrity or accuracy of the data contained herein, in particular due, but not limited to, any deletion, manipulation, or reformatting of data that may have occurred beyond its control.”

Human population and demographics were drawn from the UN WPP dataset (*World Population Prospects - Population Division - United Nations*, 2023). The KG only contains human population data from UN WPP, capturing births, deaths, total population, net migration, and more. Currently, these data are only available at the country level.

Intrinsic and extrinsic trait data for species is provided by COMBINE (Soria et al., 2021). Measured properties include litter size, body mass, home range, diet, and more for analysis of species traits that may influence infectious disease risk (B. A. Han et al., 2020).

### 2. Develop ontology

A knowledge graph depends on its structure: a semantic ontology that describes the data that need to be linked. This ontology is an abstract representation of the real-world entities, attributes, and relationships in a given domain that provides a shared vocabulary to align and integrate data from a wide range of fields (e.g., ecology, epidemiology, genomics), even if those underlying data rely upon different taxonomies and formats (see Figure 2). This domain-specific approach can offer significant value by capturing complex relationships, while also helping to build linkages between highly specialized fields.

**Figure 2.**
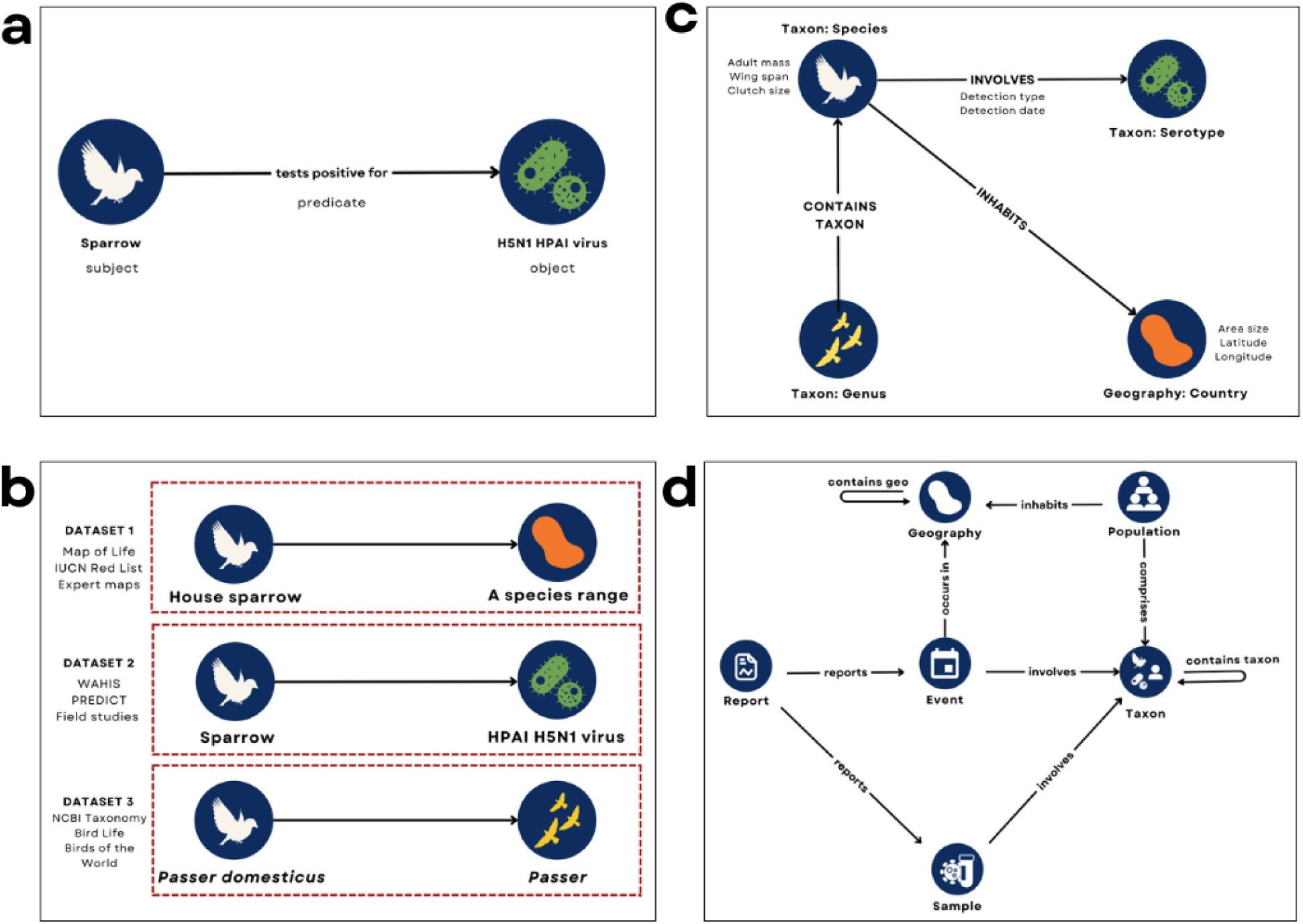
Visualizing a knowledge graph ontology. a) Representing subject-predicate-object (SPO) triples in a graph format b) The current data ecosystem; each dotted red box represents a type of tabular dataset that may contain many of the individual SPO triples. c) A domain-specific ontology to represent an outbreak. Entities are represented as blue nodes, with the bolded text above or below as labels with the node classes. To the left or right of each node are the properties of that node. d) The full ontology and schema for the zoonotic risk knowledge graph. The graph schema adheres to the ontology.

For knowledge graphs, ontologies are built around three components: nodes, edges, and properties. Nodes represent an instance of a class (derived from data, sometimes called data elements), and edges are defined as the relationships between nodes. Attributes (or properties) capture additional values or details about the node or edge. These components are semantically combined to form “triples” which describe relationships between a subject (node), a predicate (edge), and an object (another node). For instance, the statement “A sparrow tests positive for H5N1 HPAI virus” is represented with Sparrow as the subject node, H5N1 HPAI virus as the object node, and the relationship edge “tests positive for” as the predicate (Fig 2a).

The current data ecosystem is characterized by both the lack of taxonomic interoperability across datasets despite shared elements (e.g., the use of “House sparrow”, “Sparrow” and “*Passer domesticus*” to represent the same species) and the lack of information about the relationships between these entities (Fig 2b). When data are structured as columns, each column represents a node in the current data ecosystem – but there are no “labeled” relationships. Rather, the user needs to intuitively know that the relationship between the “Host” column and the “Pathogen” column is that the host “carries” the pathogen (empirically) or the host was “detected with” the pathogen (in a specific sample).

With a knowledge graph ontology, relationships are semantically labeled to precisely define the connections between entities, rather than needing to infer relationships across multiple disconnected datasets (also known as “rules mining”, (Suchanek et al., 2019). Any data referring to that species – regardless of the naming convention – would map to the same node (Fig 2c). This enables knowledge graphs to achieve interoperability and interpretability across fields without mandating the use of specific terms or measurement types in the source data. Thus, knowledge graphs with robust ontologies are a foundation for growth as new data becomes incrementally available.

While network models have a similar structure with multiple nodes and edges, all of the edges represent one kind of relationship (e.g., network models of host-pathogen associations, (Dallas et al., 2019). Knowledge graphs, on the other hand, support an infinitely scalable set of relationships to capture and analyze nuanced processes (Fig 2d).

Given the nuance of ecological relationships, including the influence of higher-order interactions, domain experts curated the taxonomy and ontology to ensure a comprehensive and accurate representation of ecological interactions and phenomena. Drawing from the 6 node classifications defined in step one, each was defined as an entity with 9 relationship types (edges) describing the zoonotic risk domain, including ecology, demographics, outbreak events, and more (Fig. 2d). The node classifications are defined in Table 1.

**Table 1.** Node classifications.

| Node type | Definition |
| --- | --- |
| Event | any ecological phenomena that occur at a time in a place, such as an outbreak |
| Report | documentation of the event, structured or unstructured |
| Taxon | any rank of a group of organisms (not individuals) from the NCBI Taxonomy |
| Geography | any coordinate location or place name/type, e.g. continent, landmark, ecological niche |
| Population | a subset of individuals of a species that occupy and interact within an area |
| Sample | any biological specimen |

**Events** are any ecological phenomena that occur at a time in a place, such as an outbreak. **Reports** are documentation of that event, structured or unstructured. Most report nodes are connected to an event; however, museum collections, archives, and passive surveillance efforts may exist only as reports with no event. For instance, in this graph, FluNet and WAHIS populate both reports and events, whereas GMPD2 populates just reports because the dataset describes known host-pathogen associations not specific to a discrete event.

**Taxon** nodes represent a rank of a group of organisms (not individuals) from the NCBI Taxonomy and are used to reconcile schematic and semantic differences arising from ecological and health-related data sources. The name of the species mentioned in a report is mapped to the TaxId with nodes for host and pathogen taxa added only when explicitly defined in a report and when host or pathogen names have a match in NCBI. We err on the side of inclusion, using the most specific name available (e.g., preserving finer classification below species level). Names that cannot be reconciled with the NCBI Taxonomy were matched to the next lowest taxonomic rank such that a report identifying Influenza A (H5) with no neuraminidase subtype is matched to Influenza A. Because some species might be either host or pathogen given some specific context, taxa were not defined by this characteristic.

Reports and events are each defined by place and time. **Geography** nodes can be as large as a continent or ecological niche and as small as an individual coordinate. These nodes are hierarchical and contain all smaller areas within them, including different types of boundaries (e.g., countries may contain states, counties, species ranges, and more). GeoNames was used to match places in the data to nodes in the KG based on name or reverse geocoded points.

Events affect **populations**, which live in primary geographies and other locations where they spend time whether to overwinter or through which they migrate. Population-specific traits are stored as metadata associated with corresponding taxon nodes.

**Sample** nodes represent data about biological specimens and laboratory analysis on those specimens and may include one or more taxa, particularly if capturing both host and pathogen species from a specific infection.

### 3. Build the knowledge graph

The process of building a knowledge graph based on the ecology-specific ontology described above is most simply described as a version of an ETL pipeline: extraction (E), transformation (T), and loading (L). ETL pipelines are the standard data engineering process of identifying, curating, and integrating datasets; knowledge graphs extend this method to facilitate interoperability between heterogeneous datasets.

In the context of knowledge graph development, extraction refers to the accessing of datasets for integration into the KG (e.g., API, remote or local .csv files) Once retrieved, we performed data cleaning which included removing incomplete rows, dropping columns not relevant to the KG, resetting data types, and splitting strings.

In this case, transformation refers to the alignment of case data to other ecological, geographic, or socioeconomic data. To support analysis across domains, host and pathogen names were matched to their taxonomic species in NCBI, and the data collection locations were standardized (e.g., using Geonames). For the former, we used NCBI e-fetch to identify host and parasite names either using taxonomic ID, if available, or fuzzy matching, where not. Each taxon was defined as a node with attributes including the taxon’s scientific name, other name synonyms, taxonomic ID, and rank. Geographic coordinates, placenames, or polygon shapefiles were then geocoded or reverse geocoded by match to their GeonamesId. This same reconciliation process for all names (e.g., host, pathogen) and places (e.g., coordinates, country names) was completed for each of the six datasets included in the knowledge graph. All other data elements were labeled to match the ontology as a node, edge, or attribute. Any unlabeled data was excluded from the KG. See Supplementary Table 1 for a list of all nodes, edges, and their properties.

Data was loaded into Neo4j graph data software using Python (3.9) and Neo4j (5.10). A series of queries were written in Cypher to reference the labels set for each data element and create the triples defined in the ontology. When run, these queries match and merge existing node patterns, and set new attributes and labels (see Supplementary File 1 for the specific queries). The data can be loaded in full at each run, or a singular source can be updated to reduce the number of times that the KG must be reconstructed. See the GitHub repository code for more detailed information (https://github.com/cghss-data-lab/uga-pipp).

### 4. Embedding: Translate the graph into matrices

Once the knowledge graph is built around the domain-specific ontology and datasets, it can be queried. While simple queries can be used for routine statistics, more complex reasoning (e.g., link prediction) requires that the information contained in knowledge graphs must be transformed into machine-readable formats. Knowledge graph embeddings are a type of machine-learning model that supports the extraction and quantification of semantic information. These embedding models traverse structural patterns in the graph to learn continuous vector representations (embeddings) of nodes and edges, which are then translated into numerical representations (matrices) (Mohamed et al., 2021; Suchanek et al., 2019). Embedding models must be iteratively trained on the data by using negative sampling to generate false triples, a scoring function to measure the likelihood of a given triple based on the patterns in the data, and/or a loss function to compare true and false triples (Lerer et al., 2019; Suchanek et al., 2019). Subsets of data are used for training to allow incremental training with quicker and more efficient adjustment of parameters.

Because knowledge graphs are relationship-centric, each type of embedding model is tailored to a particular type or direction of the relationship between nodes. We chose four embedding models each suited to different types of relationships to test which would perform best on our graph: TransE, ComplEx, DistMul, and RotatE (Wang et al., 2017). TransE (Bordes et al., 2013) is well-suited for one-to-one relationships using simple vectors, ComplEx (Trouillon et al., 2016) uses complex vectors (with real and imaginary numbers) to capture multi-dimensional relationships, DistMul (F. Yang et al., 2017) uses a bilinear diagonal model which is effective for one-to-many relationships, and RotatE (Sun et al., 2019) is adept at handling many-to-many relationships using rotational transformations

The KG embeddings were trained and optimized with the PyTorch library (*PyTorch*, n.d.) using a queried subset of the data, encompassing Geography, Taxon, and Report nodes populated from GMPD2 and FluNet. We then compared the KG embeddings against the standard metric of hits@*k*, which measures how often the models ranked the correct relationships (true triplets) within the top *k* highest-scoring (most likely) triplets (Dai et al., 2020). A higher hits@*k* score indicates better model proficiency in predicting and differentiating the connections between various entities within the KG (Dai et al., 2020; Onuki et al., 2017; Yin et al., 2018). In the biomedical domain, precision often remains below 0.2, as observed in Hetionet and BioKG, making values above this range desirable (Himmelstein et al., 2017; Lerer et al., 2019; Rivas-Barragan et al., 2022).

The results, as depicted in Figure 3a, show the precision of each model in learning the relationship types within the ontology for this graph. RotatE performed at ∼0.4 precision, indicating that it is most able to accurately predict triples, likely due to the number of many-to-many relationships within the KG (Fig. 3b). Given the structure of the graph, the strong performance of RotatE suggests that the ontology is semantically coherent, and can be used to provide reliable insights for downstream machine-learning tasks. By comparison, TransE performed the worst with less than 0.05 precision, likely because there are no one-to-one relationships represented in the ontology for it to learn (Fig. 3b). DistMul and ComplEx performed similarly; both cluster similar nodes together with accuracy, but they are unable to differentiate nodes by type (Fig. 3b).

**Figure 3a–b.**
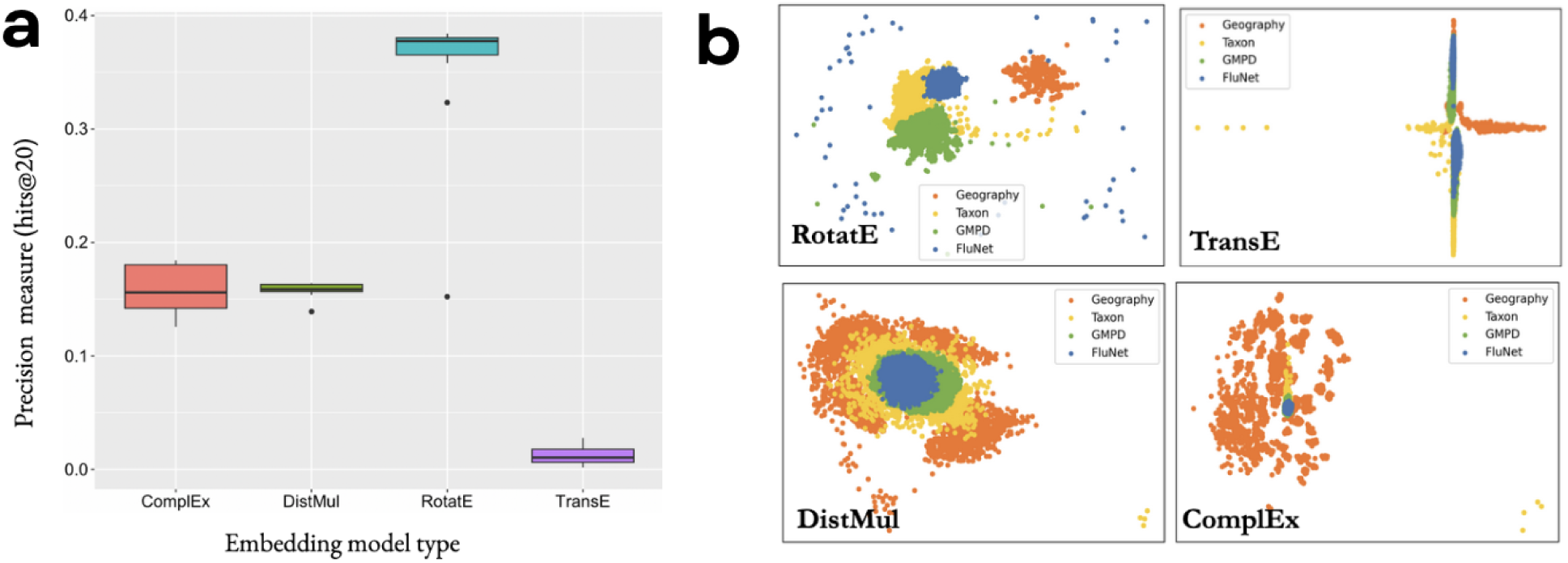
Comparison of four different KG embedding models’ performance in their ability to distinguish between node types based on connectivity patterns within the graph using data from GeoNames, NCBI Taxonomy, GMPD, and FluNet. a) Precision of KG embedding models (ComplEx, DistMul, RotatE, TransE), measured by hits@20. The y-axis indicates their precision, measured by hits@20. RotatE performed the best, and TransE performed the worst. b) KG embeddings projected in 2D space, with embedding-predicted similar nodes and relationships clustered together and dissimilar ones further apart in space. Orange dots represent nodes from GeoNames, yellow dots represent nodes from NCBI Taxonomy, green dots represent nodes from GMPD, and blue dots represent nodes from FluNet.

## Results: Applications of the KG to HPAI

The current outbreak of HPAI A/H5N1 has pandemic potential having caused unprecedented mass mortality events in wild bird populations, evolved mammal-to-mammal transmission occurring between numerous species, and human cases observed on dairy farms (Garg et al., 2024; Klaassen & Wille, 2023; Leguia et al., 2023; Sah et al., 2024). Decision-makers face the urgent need to identify when and where new cross-species transmission events are likely to occur and take immediate action to get ahead of a possible public health emergency. Challenges with gaining these insights in the status quo – as established above – are that researchers are limited to asking pairwise questions (e.g., which host carries a particular pathogen?) and that the right questions must be identified and defined before analysis. A knowledge graph, on the other hand, allows researchers to answer questions they may not even consider asking of the data, propose new hypotheses, and evaluate new areas of research. For instance, researchers could look for patterns across historical outbreaks in terms of the host species infected, populations affected, and regions impacted to determine possible characteristics of future outbreaks of that (or similar) pathogens. A knowledge graph helps visualize these connections, as well as taxonomic lineages and geographic trees, to support data exploration to suggest other relationships not captured directly in the data through inference and reasoning. This approach is useful in data-limited settings or for undersampled pathogens or host species.

One of the most important areas for future research on the reassorted HPAI A/H5 lineage involves understanding if, when, and where transmission may spread to mammal populations (Nishiura et al., 2023). Identifying mammals most heavily affected in recent and historical influenza outbreaks, and evaluating geographic patterns and characteristics of those outbreaks, may indicate relationships that can help disentangle cross-species transmission pathways and future species of concern for spillover. While there have been studies of specific populations, there has not been a global assessment of where H5 cases are occurring (Nishiura et al., 2023) in part because the data are difficult to integrate from existing sources (Klaassen & Wille, 2023) and it is traditionally not feasible to analyze patterns across numerous outbreaks given the size of the data.

We use the knowledge graph to integrate avian and mammalian case data associated with the HPAI A/H5N1 lineage for the ongoing outbreak and identify shared locations between highly affected mammals to better understand which places (and which other species within those places) may be at risk of spillover. Using a single query on data ingested into the KG (see Supplementary File 2), we find that from January 2020 to July 2023, H5N1 cases in mammals were reported in 9 countries, with over 90% of cases reported in the United States and Canada (Fig 4a). The outbreak spans 21 mammal species, with cases initially reported in *Vulpes vulpes* (Red fox) by Japan and Denmark to WAHIS on March 31, 2022, and April 1, 2022, respectively (Fig 4b). Additional H5N1 cases were reported among captive and wild *V. vulpes* in the U.S. and Canada in the following weeks. In early April, cases were reported in *Mephitis mephitis* (Striped skunk) exclusively in the U.S. and Canada (Fig 4b). These two species encompass over 70% of total H5N1 cases reported to WAHIS in mammals, which raises questions about where and how these species might interact with domesticated species or humans.

**Figures 4a–b.**
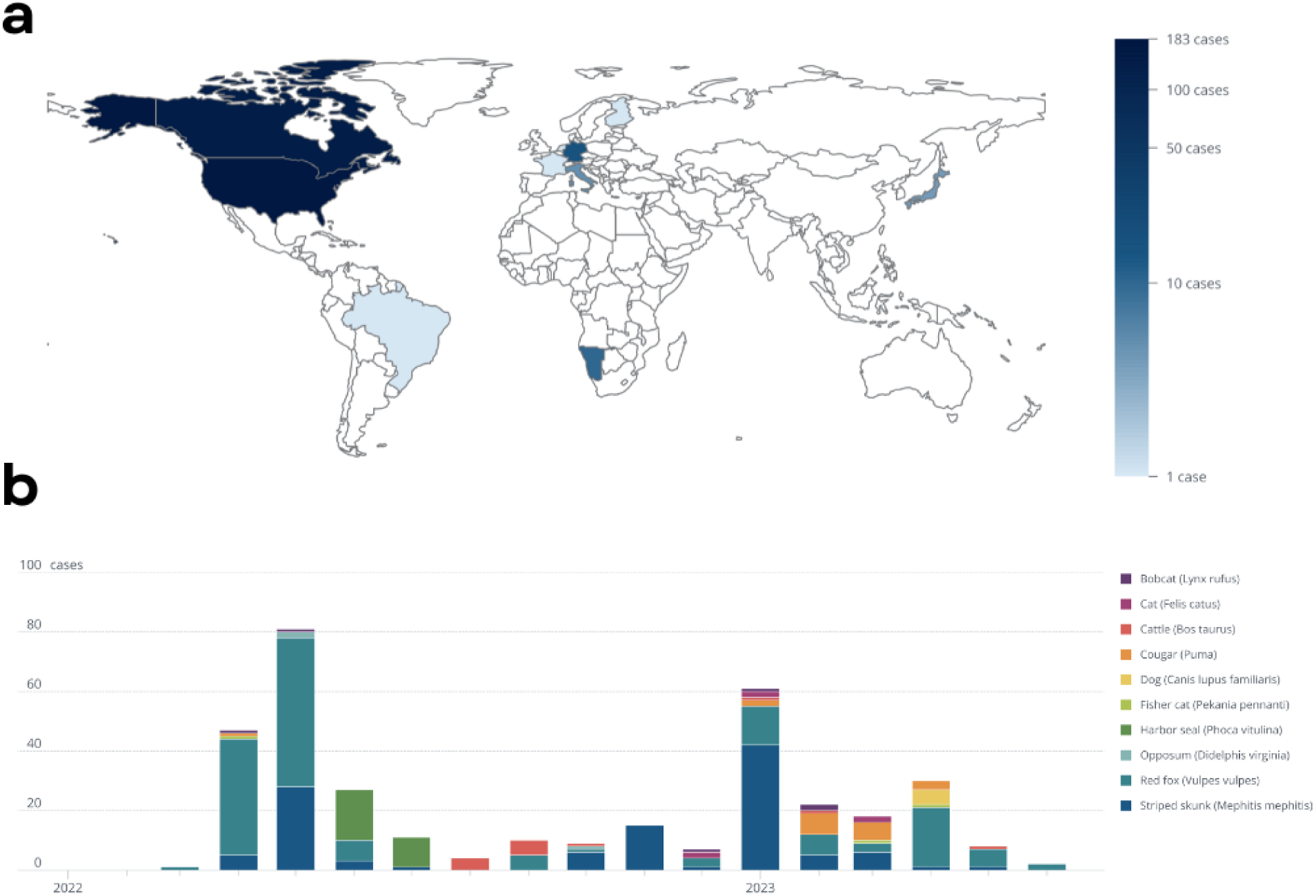
KG merges data about HPAI A/H5N1 in mammals across geographies, time, taxonomies, and scales using relationship-centric queries. a) Map of total HPAI A/H5N1 cases in mammals as reported in WAHIS from January 1, 2020 – July 1, 2023, excluding cases in humans b) Monthly counts of HPAI A/H5N1 cases from January 2020 to July 2023 for the top 10 mammal species by number of cases over time.

To expand the analysis across human and animal networks to identify places of risk and gaps in surveillance, we analyzed FluNet and WAHIS data together to explore historical influenza outbreaks involving mammals and identify locations where cross-species transmission, particularly relevant to humans, may occur based on first-order interactions (Fig 5a). Subsequently, we identified specific geographical locations that may warrant additional data collection (Fig. 5b).

**Figure 5a–b.**
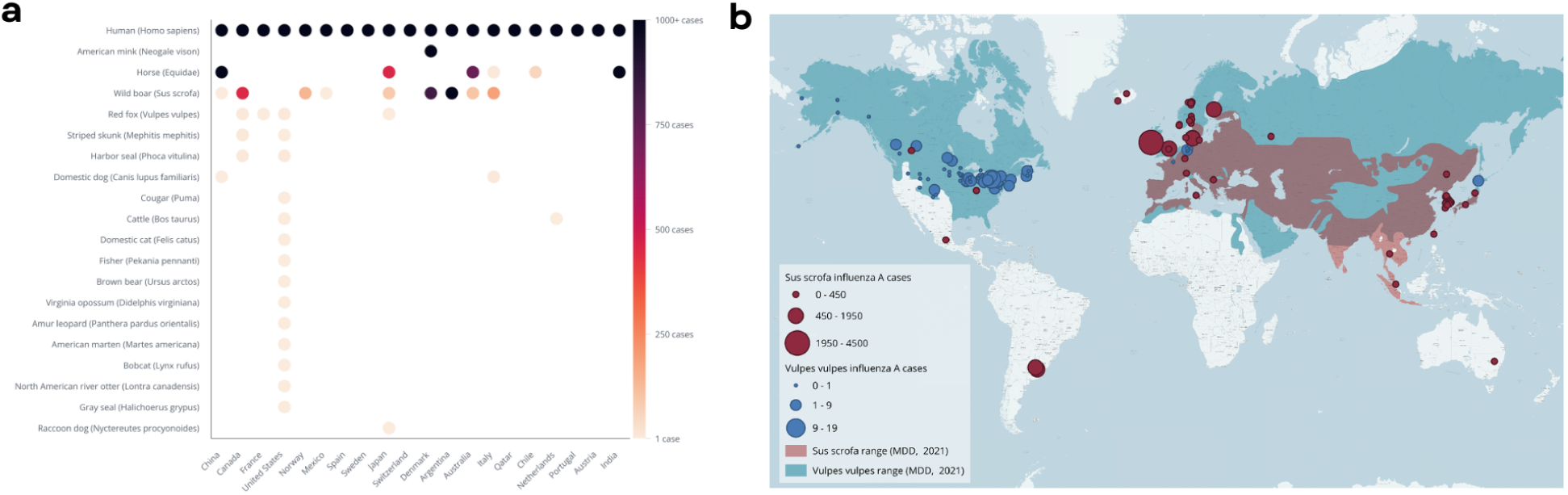
Integrating data for rapid exploratory analysis of HPAI A/H5N1. a) Heatmap showing total flu cases for all mammal species from 2005-2023, for the top 20 countries by total human cases over that time. b) Geographic distribution of cases against species range map for *Vulpes vulpes* (red fox) and *Sus scrofa* (wild boar), due to their identification as the top 2 non-domesticated species in Fig 5a.

Among the 20 countries with the highest burden of human influenza, *Sus scrofa* (boar/pig) and *V. vulpes* (red fox) were the top two mammalian species affected by influenza A (Fig. 5a). Influenza A cases in *S. scrofa* and *V. vulpes* were reported in Canada, the United States, and Japan. Notably, *S. scrofa* was not a species reported with HPAI A/H5N1, whereas *V. vulpes* had the highest number of reported cases. Analyzing the geographical range of these host species revealed significant overlap in western Europe which may be ripe for additional surveillance, especially given the proximity of these animals to humans (Fig. 5b). Countries in this region also have a high burden of human influenza. Simultaneous infections in human and animal populations that interact frequently can intensify selection pressure on the virus, creating opportunities for potential reassortment events at the animal-human interface (C. Li et al., 2010; Octaviani et al., 2010; Reperant et al., 2014).

The ease and speed of querying the knowledge graph allows for rapid evaluation of data across multiple datasets at different scales. These findings allow for rapid exploratory analysis of HPAI A/H5N1 prevalence by host. Species distribution, ecological niches, and trophic level data (from COMBINE, the IUCN Red List, Map of Life, and food web models) could be used to identify potentially susceptible or at-risk hosts. Integrating genomic data could improve the biological accuracy of host-pathogen interactions and outbreak models.

## Discussion

The KG method offers new insights more rapidly and in a more scalable manner than traditional analysis techniques. By collating the data in the KG, we were able to analyze species distribution, geospatial coverage, and temporal distribution from a single source, at a more granular level than available via the WAHIS data download alone and without needing to reconcile to new scales or taxonomies. For instance, we were able to align host and causative agent names in WAHIS (often reported as common names) to the NCBI Taxonomic backbone to facilitate linkages to other data sources and previous outbreaks involving these species. Affected places, reported by coordinate points, were rolled up to the country level by reverse geocoding and navigating through administrative levels in GeoNames.

This flexibility is in contrast to a relational or tabular format, which enforces a rigid structure that requires researchers to specify the exact columns and filtering criteria. Any data that does not fit this predefined structure is not surfaced, obscuring potentially valuable information – especially if the researcher doesn’t know to ask for it. So, while it is possible in a traditional format to produce the types of analysis performed in Figures 4 and 5, scaling up the research questions necessitates either restructuring the query over and over, re-analyzing the dataset, and seeking new data sources. These tasks are often time-consuming and labor-intensive, and may not yield comprehensive insights.

In a tabular data approach, producing the analysis shown in Figure 5 would likely involve at least 8 distinct, manual steps (see Table 2). Meanwhile, the graph-based approach streamlines the process significantly with a single query. Traversing seamlessly between geographic scales and species taxonomies, it retrieves the required data in the desired format without multiple intermediate steps. The graph handles intersections between geopolitical and natural boundaries, maintaining the integrity of point data throughout the transformation. Taxonomic integration is native to the graph, allowing for a smooth transition between common and scientific names.

**Table 2.** Steps for tabular vs. graph-based data approach.

| Tabular approach | Graph-based approach |
| --- | --- |
| Create mappings between taxonomies in different data sources, including for species and places | Mappings completed on data ingest with ontology |
| Reverse geocode coordinates from WAHIS for each influenza outbreak in mammal species, and aggregate up to the country-level |  |
| Sum all human cases of influenza (e.g. Influenza A, Influenza B, H5N1, etc.) by country from 2005-2023 | All matching data retrieved with a single query with no transformation required |
| Sum all mammal cases of influenza by mammal species and by country from 2005-2023 |  |
| Order countries by the number of human cases and select the top 20 |  |
| For each country, return the total number of mammal cases by species using the species' common name |  |
| For *Sus scrofa* and *Vulpes vulpes*, re-retrieve and merge outbreak point data |  |

Broadly, relational database management systems (RDBMS) perform best in queries requiring row-by-row access; graph databases, in contrast, perform best in queries focusing on the structure of relationships between data (Codd, 1970). As a result of these two distinct paradigms of data storage, there are differences in the types of research questions that perform best in each database type. RDBMS is the best choice when there are few many-to-many joins between datasets, there is a clear index or collection of tables that could answer the question, or the data structure itself is not of interest. While graph databases can handle many of the same questions that RDBMS can, they tend to be more difficult to work with (requiring specialized query languages) and perform slower in group by, aggregation, and sort operations. However, graph databases exhibit better performance and scalability in traversal queries, multi-table joins, and pattern matching (Patras et al., 2021). Decision trees for when to use an RDBMS or a graph database based on the research questions and structure of the data are shown in Figure 6.

**Figure 6.**
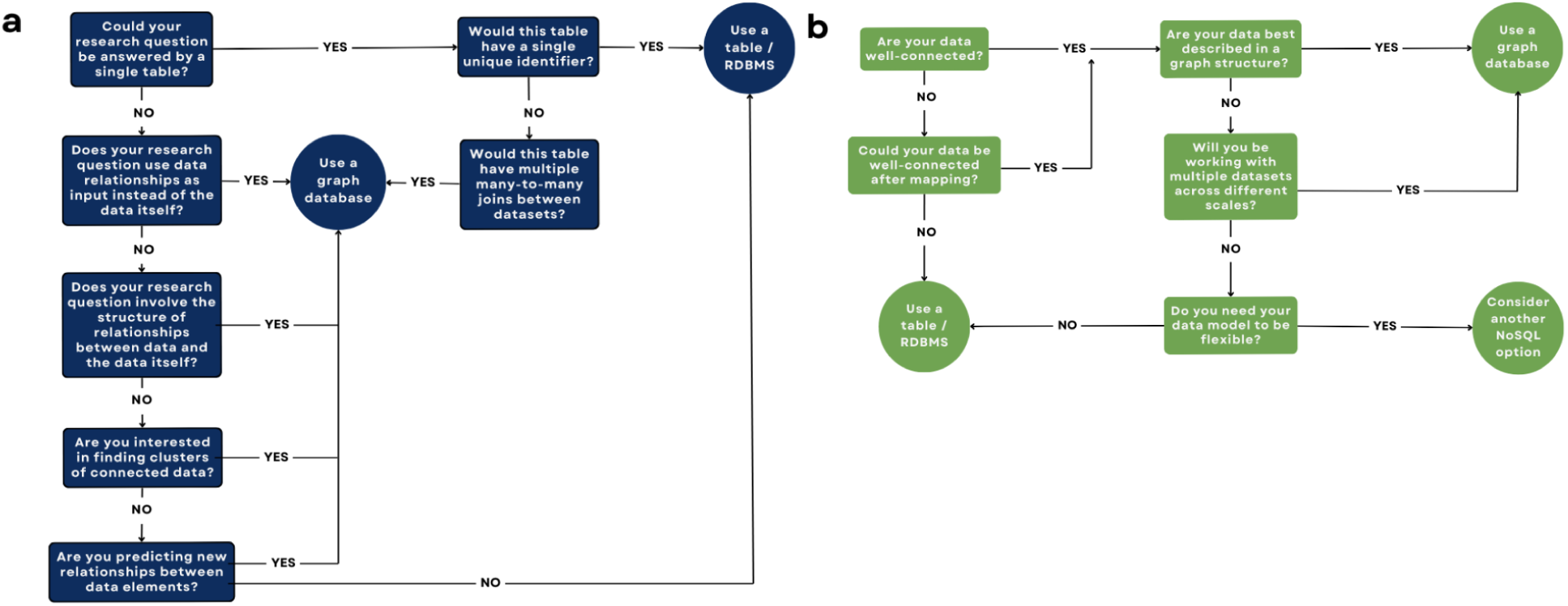
Decision tree for choosing a relational versus a graph database. a) Decision tree based on the type of research question being asked. b) Decision tree based on the structure of the data being used.

Beyond the ability to rapidly execute more complex queries for targeted results (represented by the colorful nodes in Figure 7), the knowledge graph ontology provides a cloud of interconnected data that enables deeper exploration and insight (represented by the white nodes in Figure 7). For instance, by querying for a singular host species, the graph would also surface the host’s taxonomic lineage, diet, phenotypic traits, habitat information and species range, compatible pathogens, outbreaks by which it had been affected, and any other data linked to the species. Researchers can scale their inquiries by navigating through the web of relationships in the ontology, revealing hidden connections, and adapting new research questions through interchangeable components across hundreds of hosts or pathogens. This approach proves particularly useful in response contexts, where timely and efficient data integration informs critical decisions even when sampling is limited. For example, with HPAI A/H5N1, pinpointing the next possible mammalian host is nearly impossible given how unpredictable the virus’ evolution has been (Xie et al., 2023). However, by looking for highly central host species from past outbreaks of HPAI A/H5/H7 and other influenza A virus subtypes, we can begin to ask which species should be prioritized for testing in the current outbreak to more efficiently allocate limited resources.

**Figure 7.**
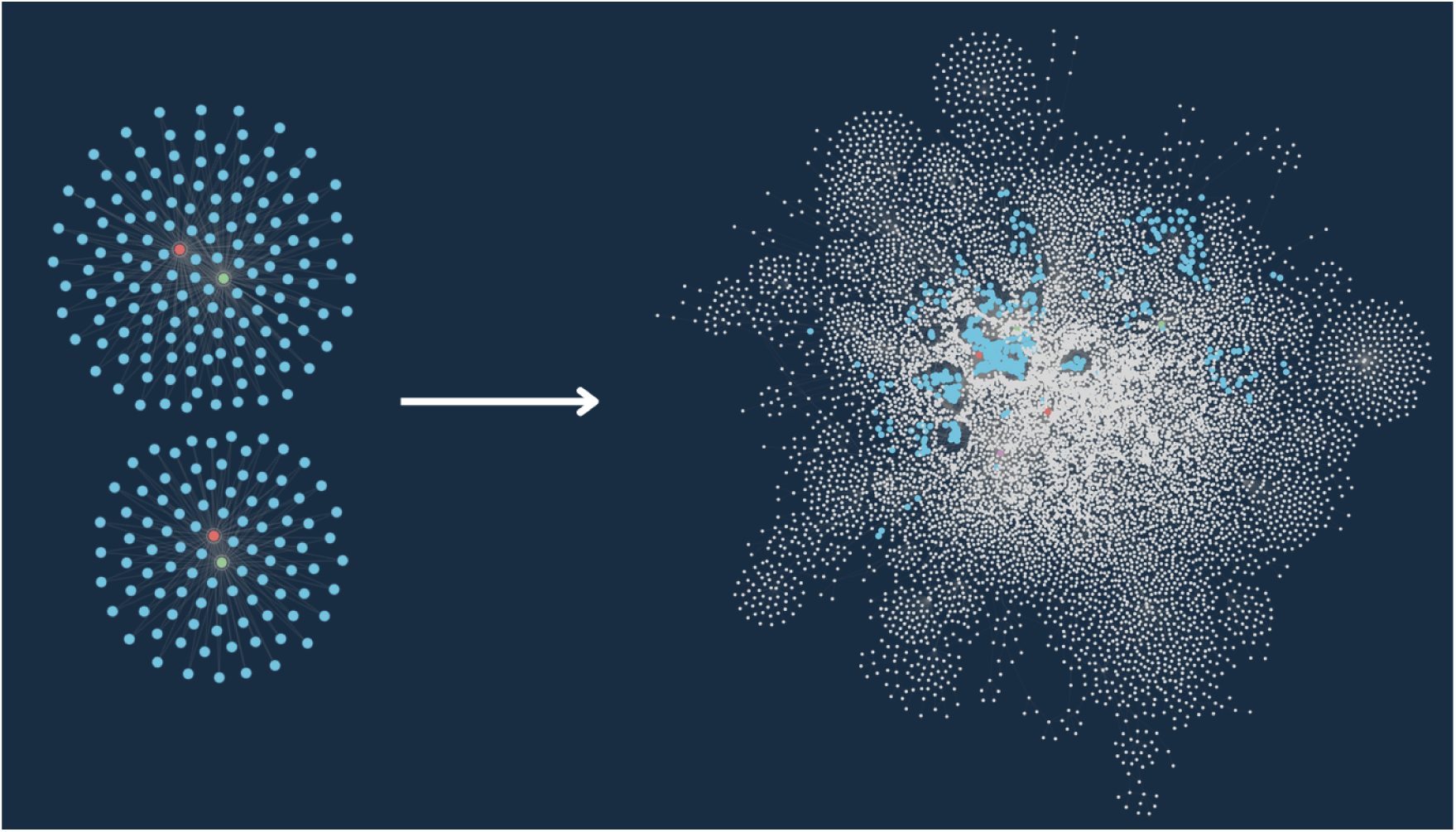
Using path expansion within the graph to surface linked data. Expanding from influenza outbreaks involving *Sus scrofa* (wild boar) and *Vulpes vulpes* (red fox) within the graph.

There are, however, several challenges to working with knowledge graphs beyond the need for technical expertise in constructing them. For instance, domains with sparse data may not be suitable for downstream modeling. When domains are inadequately represented in the knowledge graph, advanced analytical techniques have reduced pattern recognition, leading to incomplete or biased insights. Therefore, inferred relationships should be approached with skepticism and validated by subject-matter experts before being accepted as new “knowledge” to incorporate back into the graph. The development of explainable artificial intelligence may help explain the influential factors in predicting new inferred relationships and outcomes, such as disease transmission (Kelly, S., A. Ghadami, J. M. Drake, B. I. Epureanu, 2023)

The broader implications of employing knowledge graphs and ontologies for ecology – especially when combined with artificial intelligence – enable researchers to break out of traditional data silos and readily integrate expertise and data from between subfields and disciplines. As the world continues to face rapid global changes, the ability to integrate and analyze multiscale data and models will be crucial in predicting and mitigating complex systemic impacts (Pokutnaya et al., 2023). Knowledge graphs hold promise in addressing this challenge.

## Data availability

No new raw data was generated for this manuscript. Processed data is available on GitHub (https://github.com/cghss-data-lab/uga-pipp). The source code used to build the knowledge graph and generate figures 3-5 is also available on GitHub.

## Acknowledgments

This research was supported by grant DEB-2200158 from the National Science Foundation. The authors thank Justin Bahl for useful discussion and comments on previous versions of this manuscript.

## References

Bordes, A., Usunier, N., García-Durán, A., Weston, J., & Yakhnenko, O. (2013). Translating embeddings for modeling multi-relational data. Advances in Neural Information Processing Systems, 2787–2795. https://proceedings.neurips.cc/paper_files/paper/2013/file/1cecc7a77928ca8133fa24680a88d2f9-Paper.pdf

Brett, T. S., Drake, J. M., & Rohani, P. (2017). Anticipating the emergence of infectious diseases. Journal of the Royal Society, Interface / the Royal Society, 14(132). 10.1098/rsif.2017.0115

Callahan, T. J., Tripodi, I. J., Stefanski, A. L., Cappelletti, L., Taneja, S. B., Wyrwa, J. M., Casiraghi, E., Matentzoglu, N. A., Reese, J., Silverstein, J. C., Hoyt, C. T., Boyce, R. D., Malec, S. A., Unni, D. R., Joachimiak, M. P., Robinson, P. N., Mungall, C. J., Cavalleri, E., Fontana, T., … Hunter, L. E. (2024). An open source knowledge graph ecosystem for the life sciences. Scientific Data, 11(1), 363. 10.1038/s41597-024-03171-w

Chandak, P., Huang, K., & Zitnik, M. (2023). Building a knowledge graph to enable precision medicine. Scientific Data, 10(1), 67. 10.1038/s41597-023-01960-3

Codd, E. F. (1970). A relational model of data for large shared data banks. Communications of the ACM, 13(6), 377–387. 10.1145/362384.362685

Dai, Y., Wang, S., Xiong, N. N., & Guo, W. (2020). A Survey on Knowledge Graph Embedding: Approaches, Applications and Benchmarks. Electronics, 9(5), 750. 10.3390/electronics9050750

Dallas, T. A., Han, B. A., Nunn, C. L., Park, A. W., Stephens, P. R., & Drake, J. M. (2019). Host traits associated with species roles in parasite sharing networks. Oikos, 128(1), 23–32. 10.1111/oik.05602

European Food Safety Authority (EFSA), European Centre for Disease Prevention and Control (ECDC), Adlhoch, C., Alm, E., Enkirch, T., Lamb, F., Melidou, A., Willgert, K., Marangon, S., Monne, I., Stegeman, J. A., Delacourt, R., Baldinelli, F., & Broglia, A. (2024). Drivers for a pandemic due to avian influenza and options for One Health mitigation measures. EFSA Journal. European Food Safety Authority, 22(4), e8735. 10.2903/j.efsa.2024.8735

Flahault, A., Dias-Ferrao, V., Chaberty, P., Esteves, K., Valleron, A. J., & Lavanchy, D. (1998). FluNet as a tool for global monitoring of influenza on the Web. JAMA: The Journal of the American Medical Association, 280(15), 1330–1332. 10.1001/jama.280.15.1330

Friant, S. (2024). Human behaviors driving disease emergence. Evolutionary Anthropology, 33(2), e22015. 10.1002/evan.22015

Garg, S., Reed, C., Davis, C. T., Uyeki, T. M., Behravesh, C. B., Kniss, K., Budd, A., Biggerstaff, M., Adjemian, J., Barnes, J. R., Kirby, M. K., Basler, C., Szablewski, C. M., Richmond-Crum, M., Burns, E., Limbago, B., Daskalakis, D. C., Armstrong, K., Boucher, D., … Dugan, V. (2024). Outbreak of Highly Pathogenic Avian Influenza A(H5N1) Viruses in U.S. Dairy Cattle and Detection of Two Human Cases - United States, 2024. MMWR. Morbidity and Mortality Weekly Report, 73(21), 501–505. 10.15585/mmwr.mm7321e1

GeoNames. (n.d.). Retrieved July 25, 2024, from https://www.geonames.org/

Ge, X., Yang, Y., Chen, J., Li, W., Huang, Z., Zhang, W., & Peng, L. (2022). Disaster Prediction Knowledge Graph Based on Multi-Source Spatio-Temporal Information. Remote Sensing, 14(5), 1214. 10.3390/rs14051214

Gibb, R., Albery, G. F., Becker, D. J., Brierley, L., Connor, R., Dallas, T. A., Eskew, E. A., Farrell, M. J., Rasmussen, A. L., Ryan, S. J., Sweeny, A., Carlson, C. J., & Poisot, T. (2021). Data Proliferation, Reconciliation, and Synthesis in Viral Ecology. Bioscience, 71(11), 1148–1156. 10.1093/biosci/biab080

Graziosi, G., Lupini, C., Catelli, E., & Carnaccini, S. (2024). Highly Pathogenic Avian Influenza (HPAI) H5 Clade 2.3.4.4b Virus Infection in Birds and Mammals. Animals : An Open Access Journal from MDPI, 14(9). 10.3390/ani14091372

Han, B. A., & Drake, J. M. (2016). Future directions in analytics for infectious disease intelligence: Toward an integrated warning system for emerging pathogens. EMBO Reports, 17(6), 785–789. 10.15252/embr.201642534

Han, B. A., O’Regan, S. M., Paul Schmidt, J., & Drake, J. M. (2020). Integrating data mining and transmission theory in the ecology of infectious diseases. Ecology Letters, 23(8), 1178–1188. 10.1111/ele.13520

Han, F., Deng, Y., Liu, Q., Zhou, Y., Wang, J., Huang, Y., Zhang, Q., & Bian, J. (2022). Construction and application of the knowledge graph method in management of soil pollution in contaminated sites: A case study in South China. Journal of Environmental Management, 319, 115685. 10.1016/j.jenvman.2022.115685

Himmelstein, D. S., Lizee, A., Hessler, C., Brueggeman, L., Chen, S. L., Hadley, D., Green, A., Khankhanian, P., & Baranzini, S. E. (2017). Systematic integration of biomedical knowledge prioritizes drugs for repurposing. eLife, 6. 10.7554/eLife.26726

Home - Taxonomy - NCBI. (n.d.). Retrieved July 25, 2024, from https://www.ncbi.nlm.nih.gov/taxonomy

Kelly, S., A. Ghadami, J. M. Drake, B. I. Epureanu. (2023). Unveiling Key Factors in Disease Transmission through Explainable AI. MIDAS Network Annual Meeting, Atlanta, GA. Retrieved July 25, 2024, from https://www.pandemicsystems.org/DP2.html

Klaassen, M., & Wille, M. (2023). The plight and role of wild birds in the current bird flu panzootic. Nature Ecology & Evolution, 7(10), 1541–1542. 10.1038/s41559-023-02182-x

Leguia, M., Garcia-Glaessner, A., Muñoz-Saavedra, B., Juarez, D., Barrera, P., Calvo-Mac, C., Jara, J., Silva, W., Ploog, K., Amaro, Lady, Colchao-Claux, P., Johnson, C. K., Uhart, M. M., Nelson, M. I., & Lescano, J. (2023). Highly pathogenic avian influenza A (H5N1) in marine mammals and seabirds in Peru. Nature Communications, 14(1), 5489. 10.1038/s41467-023-41182-0

Lerer, A., Wu, L., Shen, J., Lacroix, T., Wehrstedt, L., Bose, A., & Peysakhovich, A. (2019). PyTorch-BigGraph: A Large-scale Graph Embedding System. In arXiv [cs.LG]. arXiv. http://arxiv.org/abs/1903.12287

Li, C., Hatta, M., Nidom, C. A., Muramoto, Y., Watanabe, S., Neumann, G., & Kawaoka, Y. (2010). Reassortment between avian H5N1 and human H3N2 influenza viruses creates hybrid viruses with substantial virulence. Proceedings of the National Academy of Sciences of the United States of America, 107(10), 4687–4692. 10.1073/pnas.0912807107

Lin, J., Zhao, Y., Huang, W., Liu, C., & Pu, H. (2021). Domain knowledge graph-based research progress of knowledge representation. Neural Computing & Applications, 33(2), 681–690. 10.1007/s00521-020-05057-5

Lin, S.-Y., Beltran-Alcrudo, D., Awada, L., Hamilton-West, C., Lavarello Schettini, A., Cáceres, P., Tizzani, P., Allepuz, A., & Casal, J. (2023). Analysing WAHIS Animal Health immediate notifications to understand global reporting trends and measure early warning capacities (2005–2021). Transboundary and Emerging Diseases, 2023, 1–10. 10.1155/2023/6666672

Li, W., Wang, S., Chen, X., Tian, Y., Gu, Z., Lopez-Carr, A., Schroeder, A., Currier, K., Schildhauer, M., & Zhu, R. (2023). GeoGraphVis: A Knowledge Graph and Geovisualization Empowered Cyberinfrastructure to Support Disaster Response and Humanitarian Aid. ISPRS International Journal of Geo-Information, 12(3), 112. 10.3390/ijgi12030112

Mohamed, S. K., Nounu, A., & Nováček, V. (2019). Drug target discovery using knowledge graph embeddings. Proceedings of the 34th ACM/SIGAPP Symposium on Applied Computing, 11–18. 10.1145/3297280.3297282

Mohamed, S. K., Nounu, A., & Nováček, V. (2021). Biological applications of knowledge graph embedding models. Briefings in Bioinformatics, 22(2), 1679–1693. 10.1093/bib/bbaa012

Nishiura, H., Kayano, T., Hayashi, K., Kobayashi, T., & Okada, Y. (2023). Letter to the Editor: Knowledge gap in assessing the risk of a human pandemic via mammals’ infection with highly pathogenic avian influenza A(H5N1). Eurosurveillance, 28(9), 2300134. 10.2807/1560-7917.ES.2023.28.9.2300134

Octaviani, C. P., Ozawa, M., Yamada, S., Goto, H., & Kawaoka, Y. (2010). High level of genetic compatibility between swine-origin H1N1 and highly pathogenic avian H5N1 influenza viruses. Journal of Virology, 84(20), 10918–10922. 10.1128/JVI.01140-10

Onuki, Y., Murata, T., Nukui, S., Inagi, S., Qiu, X., Watanabe, M., & Okamoto, H. (2017). Predicting relations of embedded RDF entities by Deep Neural Network. International Workshop on the Semantic Web. https://iswc2017.semanticweb.org/wp-content/uploads/papers/PostersDemos/paper617.pdf

Patras, V., Laskas, P., Koritsoglou, K., Fudos, I., & Karvounis, E. (2021). A comparative evaluation of RDBMS and GDBMS for shortest path operations on pedestrian navigation data. 2021 6th South-East Europe Design Automation, Computer Engineering, Computer Networks and Social Media Conference (SEEDA-CECNSM), 1–5. 10.1109/SEEDA-CECNSM53056.2021.9566235

Pestryakova, S., Vollmers, D., Sherif, M. A., Heindorf, S., Saleem, M., Moussallem, D., & Ngomo, A.-C. N. (2022). COVIDPUBGRAPH: A FAIR Knowledge Graph of COVID-19 Publications. Scientific Data, 9(1), 389. 10.1038/s41597-022-01298-2

Petersen, E., Memish, Z. A., Hui, D. S., Scagliarini, A., Simonsen, L., Simulundu, E., Bloodgood, J., Blumberg, L., Lee, S.-S., & Zumla, A. (2024). Avian “Bird” Flu – undue media panic or genuine concern for pandemic potential requiring global preparedness action? International Journal of Infectious Diseases: IJID: Official Publication of the International Society for Infectious Diseases, 145, 107062. 10.1016/j.ijid.2024.107062

Pokutnaya, D., Childers, B., Arcury-Quandt, A. E., Hochheiser, H., & Van Panhuis, W. G. (2023). An implementation framework to improve the transparency and reproducibility of computational models of infectious diseases. PLoS Computational Biology, 19(3), e1010856. 10.1371/journal.pcbi.1010856

PyTorch. (n.d.). https://pytorch.org/

Reperant, L. A., Kuiken, T., Grenfell, B. T., & Osterhaus, A. D. M. E. (2014). The immune response and within-host emergence of pandemic influenza virus. The Lancet, 384(9959), 2077–2081. 10.1016/S0140-6736(13)62425-3

Rivas-Barragan, D., Domingo-Fernández, D., Gadiya, Y., & Healey, D. (2022). Ensembles of knowledge graph embedding models improve predictions for drug discovery. Briefings in Bioinformatics, 23(6). 10.1093/bib/bbac481

Rivers, C., Chretien, J.-P., Riley, S., Pavlin, J. A., Woodward, A., Brett-Major, D., Maljkovic Berry, I., Morton, L., Jarman, R. G., Biggerstaff, M., Johansson, M. A., Reich, N. G., Meyer, D., Snyder, M. R., & Pollett, S. (2019). Using “outbreak science” to strengthen the use of models during epidemics. Nature Communications, 10(1), 3102. 10.1038/s41467-019-11067-2

Runstadler, J. A., & Puryear, W. B. (2024). The virus is out of the barn: the emergence of HPAI as a pathogen of avian and mammalian wildlife around the globe. American Journal of Veterinary Research, 85(5). 10.2460/ajvr.24.01.0018

Sah, R., Srivastava, S., Kumar, S., Mehta, R., Donovan, S., Sierra-Carrero, L., Luna, C., Woc-Colburn, L., Cardona-Ospina, J. A., Hinestroza-Jordan, M., Wessolossky, M., & Rodriguez-Morales, A. J. (2024). Concerns on H5N1 avian influenza given the outbreak in U.S. dairy cattle. Lancet Regional Health. Americas, 35, 100785. 10.1016/j.lana.2024.100785

Sánchez-Garduño, F., Castellanos, V., & Quilantán, I. (2014). Dynamics of a nonlinear mathematical model for three interacting populations. Boletín de La Sociedad Matemática Mexicana, 20(1), 147–170. 10.1007/s40590-014-0010-1

Silk, M. J., Wilber, M. Q., & Fefferman, N. H. (2022). Capturing complex interactions in disease ecology with simplicial sets. Ecology Letters, 25(10), 2217–2231. 10.1111/ele.14079

Solé, R., & Levin, S. (2022). Ecological complexity and the biosphere: the next 30 years. Philosophical Transactions of the Royal Society of London. Series B, Biological Sciences, 377(1857), 20210376. 10.1098/rstb.2021.0376

Soria, C. D., Pacifici, M., Di Marco, M., Stephen, S. M., & Rondinini, C. (2021). COMBINE: a coalesced mammal database of intrinsic and extrinsic traits. Ecology, 102(6), e03344. 10.1002/ecy.3344

Stephens, P. R., Pappalardo, P., Huang, S., Byers, J. E., Farrell, M. J., Gehman, A., Ghai, R. R., Haas, S. E., Han, B., Park, A. W., Schmidt, J. P., Altizer, S., Ezenwa, V. O., & Nunn, C. L. (2017). Global Mammal Parasite Database version 2.0. Ecology, 98(5), 1476. 10.1002/ecy.1799

Suchanek, F. M., Lajus, J., Boschin, A., & Weikum, G. (2019). Knowledge representation and rule mining in entity-centric knowledge bases. RW, 110–152. 10.1007/978-3-030-31423-1_4

Sun, Z., Deng, Z.-H., Nie, J.-Y., & Tang, J. (2019). RotatE: Knowledge Graph Embedding by Relational Rotation in Complex Space. In arXiv [cs.LG]. arXiv. http://arxiv.org/abs/1902.10197

Tan, Y., Cai, Y., Yao, R., Hu, M., & Wang, W. (2022). Complex dynamics in an eco-epidemiological model with the cost of anti-predator behaviors. Nonlinear Dynamics, 107(3), 3127–3141. 10.1007/s11071-021-07133-4

Terrestrial Code Online Access. (2021, March 23). WOAH - World Organisation for Animal Health; World Organisation for Animal Health. https://www.woah.org/en/what-we-do/standards/codes-and-manuals/terrestrial-code-online-access/

Todman, L. C., Bush, A., & Hood, A. S. C. (2023). “Small Data” for big insights in ecology. Trends in Ecology & Evolution, 38(7), 615–622. 10.1016/j.tree.2023.01.015

Trouillon, T., Welbl, J., Riedel, S., Gaussier, É., & Bouchard, G. (2016). Complex Embeddings for Simple Link Prediction. In arXiv [cs.AI]. arXiv. http://arxiv.org/abs/1606.06357

WAHIS. (n.d.). Retrieved July 25, 2024, from https://wahis.woah.org/#/event-management

Wang, Q., Mao, Z., Wang, B., & Guo, L. (2017). Knowledge Graph Embedding: A Survey of Approaches and Applications. IEEE Transactions on Knowledge and Data Engineering, 29(12), 2724–2743. 10.1109/TKDE.2017.2754499

Wardeh, M., Risley, C., McIntyre, M. K., Setzkorn, C., & Baylis, M. (2015). Database of host-pathogen and related species interactions, and their global distribution. Scientific Data, 2, 150049. 10.1038/sdata.2015.49

Woldehanna, S., & Zimicki, S. (2015). An expanded One Health model: integrating social science and One Health to inform study of the human-animal interface. Social Science & Medicine, 129, 87–95. 10.1016/j.socscimed.2014.10.059

World Animal Health Information System. (2021, March 23). WOAH - World Organisation for Animal Health; World Organisation for Animal Health. https://www.woah.org/en/what-we-do/animal-health-and-welfare/disease-data-collection/world-animal-health-information-system/

World Population Prospects - Population Division - United Nations. (2023). https://population.un.org/wpp/

Xie, R., Edwards, K. M., Wille, M., Wei, X., Wong, S.-S., Zanin, M., El-Shesheny, R., Ducatez, M., Poon, L. L. M., Kayali, G., Webby, R. J., & Dhanasekaran, V. (2023). The episodic resurgence of highly pathogenic avian influenza H5 virus. Nature, 622(7984), 810–817. 10.1038/s41586-023-06631-2

Yang, F., Yang, Z., & Cohen, W. W. (2017). Differentiable Learning of Logical Rules for Knowledge Base Reasoning. In arXiv [cs.AI]. arXiv. http://arxiv.org/abs/1702.08367

Yang, Z., Liu, C., Nie, R., Zhang, W., Zhang, L., Zhang, Z., Li, W., Liu, G., Dai, X., Zhang, D., Zhang, M., Miao, S., Fu, X., Ren, Z., & Lu, H. (2022). Research on Uncertainty of Landslide Susceptibility Prediction—Bibliometrics and Knowledge Graph Analysis. Remote Sensing, 14(16), 3879. 10.3390/rs14163879

Ye, Q., Hsieh, C.-Y., Yang, Z., Kang, Y., Chen, J., Cao, D., He, S., & Hou, T. (2021). A unified drug-target interaction prediction framework based on knowledge graph and recommendation system. Nature Communications, 12(1), 6775. 10.1038/s41467-021-27137-3

Yin, H., Zou, L., Nguyen, Q. V. H., Huang, Z., & Zhou, X. (2018). Joint Event-Partner Recommendation in Event-Based Social Networks. 2018 IEEE 34th International Conference on Data Engineering (ICDE), 929–940. 10.1109/ICDE.2018.00088

Zárate, M., Rosales, P., Braun, G., Lewis, M., Fillottrani, P. R., & Delrieux, C. (2019). OceanGraph: Some Initial Steps Toward a Oceanographic Knowledge Graph. 33–40. 10.1007/978-3-030-21395-4_3

