## Supplementary Files for "Understanding Ecological Systems Using Knowledge Graphs: An Application to Highly Pathogenic Avian Influenza"

### Supporting information

#### Supplementary Table 1: Nodes, edges, and properties

| Label | Properties | Property data source(s) |
| --- | --- | --- |
| Event (node) | Data source; Description; Start date; End date; Samples collected; Samples processed; Event ID | FluNet; WAHIS |
| Geography (node) | Data source; Place name; Feature class name; Feature code; Feature code name; Geoname ID; Latitude; Longitude | GeoNames |
| Population (node) | Data source; Report ID; Total population; Median age; Natural change in population; Population change; Births; Deaths; Life expectancy; Infant deaths; Under five deaths; Net migration | UN World Pop |
| Report (node) | Data source; Reason for report; Report date; Report description; Report ID | FluNet; WAHIS |
| Sample (node) | Data source; Sample ID; NCBI Accession ID; Collection date; Collection institution | GMPD-2 |
| Taxon (node) | Data source; Name; Rank; Tax ID; Adult mass; Brain mass; Adult body length; Adult forearm length; Maximum longevity; Days to maturity; Female days to maturity; Male days to maturity; Days to first reproductive cycle; Gestation length; Teat number; Litter size; Litters per year; Interbirth interval; Neonate mass; | NCBI Taxonomy; COMBINE |

| | | |
|---|---|---|
| | Weaning age; Weaning mass; Generation length; Dispersal; Density; Home range; Social group size; Percent of diet from invertebrates; Percent of diet from vertebrates; Percent of diet from plants; Percent of diet from mammals and birds; Percent of diet from reptiles and amphibians; Percent of diet from fish; Percent of diet from unknown vertebrates; Percent of diet from dead organisms; Percent of diet from fruits; Percent of diet from nectar and pollen; Percent of diet from seeds; Percent of diet from other plant materials; Number of dietary categories in diet; Upper elevation boundary; Lower elevation boundary; Altitude breadth; Habitat breadth; Hibernation; Freshwater habitat; Marine habitat; Terrestrial non-volant habitat; Terrestrial volant habitat; Island dwelling; Dissected by mountains; Glaciation; Fossoriality source; Trophic level; Foraging stratum; Activity cycle; Island endemicity | |
| CONTAINS_GEO (edge) | N/A | |
| CONTAINS_TAX (edge) | N/A | |
| INVOLVES (edge) | Observation date; Observation type; Positive cases; Samples collected; Samples processed; Subtype; Deaths; Role; Wild species | GMPD-2; FluNet; WAHIS |
| OCCURS_IN (edge) | N/A | |
| REPORTS (edge) | N/A | |

#### Supplementary File 1: Queries for ingest

*COMBINE:*

```
UNWIND $Mapping as mapping
MERGE (tax:Taxon {name : mapping.iucn2020_binomial})
ON MATCH SET
      tax.adult_mass_g = toFloat(mapping.adult_mass_g),
      tax.brain_mass_g = toFloat(mapping.brain_mass_g),
      tax.adult_body_length_mm = toFloat(mapping.adult_body_length_mm),
      tax.adult_forearm_length_mm = toFloat(mapping.adult_forearm_length_mm),
      tax.max_longevity_d = toFloat(mapping.max_longevity_d),
      tax.maturity_d = toFloat(mapping.maturity_d),
      tax.female_maturity_d = toFloat(mapping.female_maturity_d),
      tax.male_maturity_d = toFloat(mapping.male_maturity_d),
      tax.age_first_reproduction_d = toFloat(mapping.age_first_reproduction_d),
      tax.gestation_length_d = toFloat(mapping.gestation_length_d),
      tax.teat_number_n = toFloat(mapping.teat_number_n),
      tax.litter_size_n = toFloat(mapping.litter_size_n),
      tax.litters_per_year_n = toFloat(mapping.litters_per_year_n),
      tax.interbirth_interval_d = toFloat(mapping.interbirth_interval_d),
      tax.neonate_mass_g = toFloat(mapping.neonate_mass_g),
      tax.weaning_age_d = toFloat(mapping.weaning_age_d),
      tax.weaning_mass_g = toFloat(mapping.weaning_mass_g),
      tax.generation_length_d = toFloat(mapping.generation_length_d),
      tax.dispersal_km = toFloat(mapping.dispersal_km),
      tax.density_n_km2 = toFloat(mapping.density_n_km2),
      tax.home_range_km2 = toFloat(mapping.home_range_km2),
      tax.social_group_n = toFloat(mapping.social_group_n),
      tax.dphy_invertebrate = toFloat(mapping.dphy_invertebrate),
      tax.dphy_vertebrate = toFloat(mapping.dphy_vertebrate),
      tax.dphy_plant = toFloat(mapping.dphy_plant),
      tax.det_inv = toFloat(mapping.det_inv),
      tax.det_vend = toFloat(mapping.det_vend),
      tax.det_vect = toFloat(mapping.det_vect),
      tax.det_vfish = toFloat(mapping.det_vfish),
      tax.det_vunk = toFloat(mapping.det_vunk),
      tax.det_scav = toFloat(mapping.det_scav),
      tax.det_fruit = toFloat(mapping.det_fruit),
      tax.det_nect = toFloat(mapping.det_nect),
      tax.det_seed = toFloat(mapping.det_seed),
      tax.det_plantother = toFloat(mapping.det_plantother),
      tax.det_diet_breadth_n = toFloat(mapping.det_diet_breadth_n),
      tax.upper_elevation_m = toFloat(mapping.upper_elevation_m),
      tax.lower_elevation_m = toFloat(mapping.lower_elevation_m),
      tax.altitude_breadth_m = toFloat(mapping.altitude_breadth_m),
      tax.habitat_breadth_n = toFloat(mapping.habitat_breadth_n),
      tax.hibernation_torpor = mapping.hibernation_torpor,
      tax.freshwater = mapping.freshwater,
      tax.marine = mapping.marine,
      tax.terrestrial_non_volant = mapping.terrestrial_non_volant,
      tax.terrestrial_volant = mapping.terrestrial_volant,
      tax.island_dwelling = mapping.island_dwelling,
      tax.disected_by_mountains = mapping.disected_by_mountains,
      tax.glaciation = mapping.glaciation,
      tax.fossoriality   = mapping.fossoriality,
      tax.trophic_level  = mapping.trophic_level,
```

```
      tax.foraging_stratum  = mapping.foraging_stratum,
      tax.activity_cycle   = mapping.activity_cycle,
      tax.island_endemicity = mapping.island_endemicity
FOREACH (realm in mapping.biogeographical_realm |
      MERGE (geo:Geography {name : realm, data_source : "COMBINE" })
      MERGE (tax)-[:INHABITS]->(geo)
)
```

*FluNet:*

```
UNWIND $Mapping AS mapping
CREATE (flunet:Report {report_id : toInteger(mapping.report_id), data_source:
'FluNet'})
CREATE (event:Event {event_id : mapping.eventId,
      data_source : 'FluNet',
      start_date : DATE(mapping.start_date),
      end_date : DATE(mapping.end_date),
      collected : toFloat(mapping.Collected),
      processed : toFloat(mapping.Processed)})

MERGE (host:Taxon {tax_id : toInteger(9606),
      name : 'Homo sapiens',
      rank : 'species',
      data_source : 'NCBI Taxonomy'})
MERGE (influenzaA:Taxon {tax_id : toInteger(11320),
      name : "Influenza A virus",
      rank : "species",
      data_source : 'NCBI Taxonomy'})
MERGE (influenzaB:Taxon {tax_id : toInteger(11520),
      name : "Influenza B virus",
      rank : "species",
      data_source : 'NCBI Taxonomy'})
MERGE (influenzaH:Taxon {tax_id : toInteger(114727),
      name : "H1N1 subtype",
      rank : "serotype",
      data_source : 'NCBI Taxonomy'})

MERGE (flunet)-[:REPORTS]->(event)
MERGE (event)-[:INVOLVES {role : 'host'}]->(host)

FOREACH (map in (CASE WHEN mapping.AH1 <> '' THEN [1] ELSE [] END) |
      MERGE (event)-[:INVOLVES {role : 'pathogen', subtype : 'A(H1)',
      positive : toFloat(mapping.AH1), deaths: 'NA', observation_type :
"Laboratory detection"}]->(influenzaA))

FOREACH (map in (CASE WHEN mapping.AH1N1 <> '' THEN [1] ELSE [] END) |
      MERGE (event)-[:INVOLVES {role : 'pathogen',
      positive : toFloat(mapping.AH1N1), deaths: 'NA', observation_type :
"Laboratory detection"}]->(influenzaH))

FOREACH (map in (CASE WHEN mapping.AH3 <> '' THEN [1] ELSE [] END) |
      MERGE (event)-[:INVOLVES {role : 'pathogen', subtype : 'A(H3)',
      positive : toFloat(mapping.AH3), deaths: 'NA', observation_type :
"Laboratory detection"}]->(influenzaA))

FOREACH (map in (CASE WHEN mapping.AH5 <> '' THEN [1] ELSE [] END) |
      MERGE (event)-[:INVOLVES {role : 'pathogen', subtype : 'A(H5)',
```

```
	positive : toFloat(mapping.AH5), deaths: 'NA', observation_type :
"Laboratory detection"}]->(influenzaA))

FOREACH (map in (CASE WHEN mapping.Anotsubtyped <> '' THEN [1] ELSE [] END) |
	MERGE (event)-[:INVOLVES {role : 'pathogen', subtype : 'NA',
	positive : toFloat(mapping.Anotsubtyped), deaths: 'NA', observation_type
: "Laboratory detection"}]->(influenzaA))

FOREACH (map in (CASE WHEN mapping.BYamagata <> '' THEN [1] ELSE [] END) |
	MERGE (event)-[:INVOLVES {role : 'pathogen', subtype : 'Yamagata',
	positive : toFloat(mapping.BYamagata), deaths: 'NA', observation_type :
"Laboratory detection"}]->(influenzaB))

FOREACH (map in (CASE WHEN mapping.BVictoria <> '' THEN [1] ELSE [] END) |
	MERGE (event)-[:INVOLVES {role : 'pathogen', subtype : 'Victoria',
	positive : toFloat(mapping.BVictoria), deaths: 'NA', observation_type :
"Laboratory detection"}]->(influenzaB))

FOREACH (map in (CASE WHEN mapping.Bnotsubtyped <> '' THEN [1] ELSE [] END) |
	MERGE (event)-[:INVOLVES {role : 'pathogen', subtype : 'NA',
	positive : toFloat(mapping.Bnotsubtyped), deaths: 'NA', observation_type
: "Laboratory detection"}]->(influenzaB))

FOREACH (map in (CASE WHEN mapping.geonames.geonameId IS NOT NULL THEN [1] ELSE
[] END) |
	MERGE (territory:Geography {geoname_id : mapping.geonames.geonameId})
	ON CREATE SET
	territory.data_source = 'GeoNames',
	territory.geoname_id = toInteger(mapping.geonames.geonameId),
	territory.name = mapping.geonames.name,
	territory.admin_type = mapping.geonames.adminType,
	territory.iso2 = mapping.geonames.iso2,
	territory.fcl_name = mapping.geonames.fclName,
	territory.fcode_name = mapping.geonames.fcodeName,
	territory.lat = toFloat(mapping.geonames.lat),
	territory.long = toFloat(mapping.geonames.lng),
	territory.fcode = mapping.geonames.fcode
	MERGE (event)-[:OCCURS_IN]->(territory))
```

*GeoNames:*

```
UNWIND $Mapping as mapping
MERGE (geo:Geography {geoname_id : toInteger(mapping.geonameId)})
ON CREATE SET
	geo.name = mapping.name,
	geo.admin_type = mapping.adminType,
	geo.iso2 = mapping.iso2,
	geo.fcl_name = mapping.fclName,
	geo.fcode_name = mapping.fcodeName,
	geo.lat = toFloat(mapping.lat),
	geo.long = toFloat(mapping.lng),
	geo.fcode = mapping.fcode
WITH COLLECT(geo) AS hierarchy
UNWIND RANGE(0, SIZE(hierarchy) - 2) as idx
WITH hierarchy[idx] AS h1, hierarchy[idx+1] AS h2
MERGE (h1)-[:CONTAINS_GEO]->(h2)
```

*GMPD-2:*

```
UNWIND $Mapping AS mapping
CREATE (gmpd:Report {data_source: "GMPD",
                     reference : mapping.Citation,
                     report_id: toInteger(mapping.report_id)})

CREATE (sample:Sample {data_source: "GMPD", processed:
toFloat(mapping.processed)})

MERGE (gmpd)-[:REPORTS]->(sample)
// Process host information
FOREACH (map in (CASE WHEN mapping.Host.taxId IS NOT NULL THEN [1] ELSE [] END)
|
      MERGE (host:Taxon {tax_id : toInteger(mapping.Host.taxId)})
      ON CREATE SET
      host.name = mapping.Host.name,
      host.rank = mapping.Host.rank,
      host.data_source = "NCBI Taxonomy"
      MERGE (sample)-[:INVOLVES {role : 'host'}]->(host))

// Process pathogen information
FOREACH (map in (CASE WHEN mapping.Parasite.taxId IS NOT NULL THEN [1] ELSE []
END) |
      MERGE (pathogen:Taxon {tax_id : toInteger(mapping.Parasite.taxId)})
      ON CREATE SET
      pathogen.name = mapping.Parasite.name,
      pathogen.rank = mapping.Parasite.rank,
      pathogen.data_source = "NCBI Taxonomy"
      MERGE (sample)-[:INVOLVES {role : 'pathogen',
      observation_type : mapping.SamplingType,
      positive : toFloat(mapping.positive),
      deaths : "NA",
      species_wild : toBoolean(mapping.species_wild)
      }]->(pathogen))

// Process geographical location
FOREACH (map in (CASE WHEN mapping.location.geonameId IS NOT NULL THEN [1] ELSE
[] END) |
      MERGE (territory:Geography {geoname_id :
toInteger(mapping.location.geonameId)})
      ON CREATE SET
      territory.data_source = 'GeoNames',
      territory.geoname_id = toInteger(mapping.location.geonameId),
      territory.name = mapping.location.name,
      territory.admin_type = mapping.location.adminType,
      territory.iso2 = mapping.location.iso2,
      territory.fcl_name = mapping.location.fclName,
      territory.fcode_name = mapping.location.fcodeName,
      territory.lat = toFloat(mapping.location.lat),
      territory.long = toFloat(mapping.location.lng),
      territory.fcode = mapping.location.fcode
      MERGE (sample)-[:OCCURS_IN]->(territory))
```

*NCBI Taxonomy:*

```
UNWIND $Mapping as mapping
MERGE (tax:Taxon {tax_id : toInteger(mapping.taxId)})
ON CREATE SET
      tax.name = mapping.name,
      tax.rank = mapping.rank,
      tax.data_source = mapping.data_source
WITH COLLECT(tax) AS hierarchy
UNWIND RANGE(0, SIZE(hierarchy) - 2) as idx
WITH hierarchy[idx] AS h1, hierarchy[idx+1] AS h2
MERGE (h1)-[:CONTAINS_TAX]->(h2)
```

*UN World Pop:*

```
UNWIND $Mapping AS mapping
CREATE (population:Population {
      data_source : mapping.data_source,
      report_id : mapping.report_id,
      date : DATE(mapping.date),
      total_population : toFloat(mapping.TPopulation1July),
      median_age : toFloat(mapping.MedianAgePop),
      natural_change : toFloat(mapping.NatChange),
      population_change : toFloat(mapping.PopChange),
      births : toFloat(mapping.Births),
      deaths : toFloat(mapping.Deaths),
      life_expectancy : toFloat(mapping.LEx),
      infant_deaths : toFloat(mapping.InfantDeaths),
      under_five_deaths : toFloat(mapping.Under5Deaths),
      net_migration : toFloat(mapping.NetMigrations)})
MERGE (geography:Geography {geoname_id: toInteger(mapping.geonames.geonameId)})
MERGE (population)-[:INHABITS]->(geography)
```

*WAHIS:*

```
UNWIND $Mapping AS mapping
MERGE (report:Report {report_id: toInteger(mapping.report.reportId)})
ON CREATE SET
      report.data_source = "WAHIS",
      report.report_date = DATE(mapping.report.reportedOn),
      report.reason_for_report = mapping.event.reason.translation,
      report.report_description = mapping.event.eventComment

MERGE (event:Event {event_id : mapping.outbreak.outbreakId})
ON CREATE SET
      event.start_date = DATE(mapping.outbreak.start_date),
      event.end_date = DATE(mapping.outbreak.end_date),
      event.description = mapping.outbreak.description,
      event.data_source = "WAHIS"

MERGE (report)-[:REPORTS]->(event)

// Set geographical information
```

```
MERGE (territory:Geography {geoname_id :
toInteger(mapping.outbreak.geonames.geonameId)})
ON CREATE SET
      territory.data_source = 'GeoNames',
      territory.geoname_id = toInteger(mapping.outbreak.geonames.geonameId),
      territory.name = mapping.outbreak.geonames.name,
      territory.admin_type = mapping.outbreak.geonames.adminType,
      territory.iso2 = mapping.outbreak.geonames.iso2,
      territory.fcl_name = mapping.outbreak.geonames.fclName,
      territory.fcode_name = mapping.outbreak.geonames.fcodeName,
      territory.lat = toFloat(mapping.outbreak.geonames.lat),
      territory.long = toFloat(mapping.outbreak.geonames.lng),
      territory.fcode = mapping.outbreak.geonames.fcode
MERGE (event)-[:OCCURS_IN]->(territory)

// Set host information (mapping.hosts = array of dictionaries)
// Skip host data list if tax ID is null
FOREACH (hostDataList in mapping.hosts |
      FOREACH (hostData in (CASE WHEN hostDataList.taxId IS NOT NULL THEN
hostDataList ELSE [] END) |
            MERGE (host:Taxon {tax_id: toInteger(hostData.taxId)})
            ON CREATE SET
                        host.name = hostData.name,
                        host.rank = hostData.rank,
                        host.data_source = "NCBI Taxonomy"
            MERGE (event)-[involves:INVOLVES {role: 'host'}]->(host)
            ON CREATE SET
                        event.processed = toFloat(hostData.processed),
                        involves.positive = toFloat(hostData.positive),
                        involves.deaths = toFloat(hostData.deaths),
                        involves.observation_type = hostData.observation_type,
                        involves.observation_date =
DATE(hostData.observation_date),
                        involves.species_wild =
toBoolean(hostData.species_wild)
      ))

// Process pathogen information
FOREACH (map in (CASE WHEN mapping.pathogen.taxId IS NOT NULL THEN [1] ELSE []
END) |
      MERGE (pathogen:Taxon {tax_id : toInteger(mapping.pathogen.taxId)})
      ON CREATE SET
            pathogen.name = mapping.pathogen.name,
            pathogen.rank = mapping.pathogen.rank,
            pathogen.data_source = "NCBI Taxonomy"
      MERGE (event)-[:INVOLVES {role : 'pathogen'} ]->(pathogen))
```

#### Supplementary File 2: Queries for analysis

```
// HPAI cases by month, country, and mammal species
MATCH (hpai:Taxon)
WHERE hpai.name STARTS WITH "H5" OR hpai.name STARTS WITH "H7"
MATCH (t:Taxon {name:"Mammalia"})-[c:CONTAINS_TAX*]->(mammals:Taxon)
MATCH (hpai)<-[inv:INVOLVES {role:
'pathogen'}]-(event:Event)-[involves:INVOLVES
{role:'host'}]->(mammals),
      (event)-[o:OCCURS_IN]->(place:Geography)
WHERE DATE(event.start_date) >= DATE("2020-01-01") AND
DATE(event.start_date) <= DATE("2023-07-01")
MATCH (place)<-[:CONTAINS_GEO*]-(country:Geography {fcode:"PCLI"})
WITH mammals.name AS mammalName, country.name AS countryName,
apoc.temporal.format(event.start_date, 'yyyy-MM') AS month,
sum(toInteger(involves.positive)) AS totalCases
WHERE totalCases > 0
RETURN mammalName AS species, countryName AS country, month AS
yearMonth, totalCases AS monthlyCases
ORDER BY month ASC

// All influenza A cases by date, place, host species, host class,
and pathogen subtype
MATCH (parent:Taxon {name: 'Influenza A
virus'})-[:CONTAINS_TAX*0..]->(virus:Taxon)
OPTIONAL MATCH (virus)<-[:INVOLVES {role:
'pathogen'}]-(event:Event)-[involves:INVOLVES {role:
'host'}]->(species:Taxon), (event)-[:OCCURS_IN]->(g:Geography)
OPTIONAL MATCH (species)<-[:CONTAINS_TAX*]-(t:Taxon {rank: "Class"})
WHERE involves.positive IS NOT NULL AND involves.positive <> 'NA' AND
species.name IS NOT NULL
WITH g.name AS place, g.lat as lat, g.long as long, event.start_date
AS date, SUM(toInteger(involves.positive)) AS totalCases, t.name AS
class, species.name as host, parent.name as species, virus.name as
subtype
RETURN place, lat, long, date, class, host, totalCases, species,
subtype
ORDER BY date DESC
```
